# Translation efficiency changes at heat shock in *Saccharomyces cerevisiae*

**DOI:** 10.64898/2025.12.09.693274

**Authors:** Rodolfo L. Carneiro, Tatiana Domitrovic, Fernando L. Palhano

## Abstract

Both eukaryotic and prokaryotic cells respond to heat shock by engaging multiple regulatory mechanisms that preserve proteostasis, including adjustments in transcription, translation, and mRNA metabolism. Although the transcriptional response to heat stress in *Saccharomyces cerevisiae* has been well characterized, the extent to which translation efficiency (TE) is remodeled and how this remodeling contributes to protein synthesis remains less understood. Here, analysis of previously published ribosome profiling and RNA-seq data revealed that in *S. cerevisiae*, the TE varies during heat shock, depending on both temperature and stress duration. In contrast, TE remained largely stable in *Escherichia coli* under comparable conditions or during activation of σ32, the main regulator of heat shock response in bacteria. In yeast, TE modulation patterns correlated with changes in protein abundance and differed markedly between early and late stages of stress. At 10 min, transcripts with short 5′UTRs and high codon optimality tended to display higher TE, primarily due to the easier ribosome engagement in these transcripts. By 30 min, the TE pattern across transcripts had changed drastically, suggesting increased availability of newly synthesized stress-response transcripts. These findings support a model in which early TE adjustments depend on intrinsic mRNA properties, whereas shifts in mRNA accessibility and transcriptional output shape later changes. The results highlight TE remodeling as a dynamic component of the heat-shock response in yeast and distinguish this behavior from the more buffered response observed in bacteria.

## Introduction

Exposure to elevated temperature disrupts protein folding and challenges the cellular systems responsible for maintaining proteostasis (**Ghadanian et al, 2025**). To counter these effects, cells activate a conserved heat-shock response that includes rapid induction of molecular chaperones, repression of housekeeping functions, and formation of stress-induced ribonucleoprotein assemblies. In addition to transcriptional remodeling, translational control plays a central role in determining which proteins are synthesized during stress (**Desroches Altamirano and Alberti, 2025**). Translation efficiency (TE), defined as the ratio of ribosome occupancy to mRNA abundance, reflects how extensively ribosomes are translating each molecule of mRNA. It suggests that regulatory mechanisms at the translational level, rather than RNA abundance, mediate transcriptional regulation. TE varies substantially across genes and can be influenced by 5 ′ UTR length, secondary structure, codon usage, and other sequence-dependent features (**Razumova et al., 2025**). In bacteria, TE is known to be relatively stable during heat shock, even when the response is driven by alternative sigma factors or stabilized variants (**Morgan et al, 2018**). Whether eukaryotic systems exhibit a similar buffering capacity remains unclear.

Recent studies have shown that heat shock in yeast induces alterations in tRNA pools, promotes condensation of RNA-binding proteins and mRNAs, and changes the activity of translation initiation factors (**Verghese et al, 2012, Torrent et al., 2018, Desroches Altamirano and Alberti, 2025)**. These processes suggest that the translational response to heat shock may involve mechanisms distinct from those described in bacteria and could ultimately lead to TE changes as a regulatory mechanism.

In this study, we examined previously published datasets on TE and tRNA availability under different temperature and exposure time conditions to quantify how TE changes during heat shock in *Saccharomyces cerevisiae* and to identify features associated with increased or decreased translation. By comparing yeast and bacterial responses, and by examining both early and prolonged stress conditions, we aimed to understand how translational regulation contributes to the reorganization of protein synthesis during heat shock.

## Methods

### Data Sources

Translation efficiency, ribosome profiling, mRNA-seq, and proteomics data from yeast heat-shock genes were obtained from **Mühlhofer et al., 2019**. Translation efficiency of yeast exposed to hydrogen peroxide was obtained from **Gerashchenko et al, 2012**. Bacterial translation efficiency was obtained from **Morgan et al. (2018)**.

The stress-adjusted tAI (s.tAI) was obtained from **Torrent et al., 2018**.

The amount of optimal codons was obtained from *the Saccharomyces* Genome Database. The 5 ′ UTR length was obtained from **Zhan et al., 2025**.

### Samples definition

In Figure 2, each data point shown in the histogram and box-plot panels corresponds to a gene for which translational efficiency (TE) values were available under both conditions (25ºC and heat shock), as well as the corresponding protein expression fold change. In Figures 3, 4, S1, and S2, each data point represents a gene for which TE values were obtained for both conditions displayed in the respective figure.

### Statistical analyses and raw data

The raw data used to create all figures are in Table S1. All statistical analyses were performed with GraphPad Prism 7. A nonparametric Kolmogorov-Smirnov t-test was used in Figure 2, where **** p value < 0.0001, ** p value = 0.0039. For Figures 3, 4, S1, and S2, a nonparametric ANOVA Kruskal-Wallis test with a control sample as reference was used: **** p value < 0.0001, *** p value < 0.001, ** p value < 0.01.

## Results and discussion

First, we validated whether ribosome profiling analysis corroborates the previously described changes in translation efficiency (TE) in *Saccharomyces cerevisiae* under heat shock. When TE values measured at 25 °C were compared with those collected after exposure to 37 °C or 42 °C (**Mühlhofer et al, 2019**), the Pearson correlation values spanned a broad range (0.57 to 0.85), indicating substantial reorganization of translational activity under these conditions, if compared with *Escherichia coli* (**Fig. 1A**). A similar extent of variability was observed during oxidative stress (**Fig. 1B**) (**Gerashchenko et al, 2012**). In contrast, TE remained highly consistent in *Escherichia coli* when cells were shifted to 42 °C. TE also remained unchanged when the key regulator of the bacterial heat-shock response, σ32, or its stabilized variant, σ32-I54N, was expressed from a plasmid. (**Fig. 1C**) (**Morgan et al, 2018**). These comparisons show that TE is strongly modulated by temperature stress in yeast, whereas in bacteria it is not. To explore how these translational changes relate to the proteome in *S. cerevisiae*, we integrated TE measurements derived from ribosome profiling datasets with proteomic data obtained after 30 min at 37 °C (**Mühlhofer et al, 2019**). Protein abundance ratios (37 °C/25 °C) grouped into distinct patterns (**Fig. 2A**), and when TE values were examined under matching conditions, transcripts associated with reduced protein levels tended to exhibit lower TE at 37 °C. Likewise, those associated with increased protein levels showed higher TE values (**Fig. 2B–D**). Equivalent behavior was observed when heat shock was performed at 42 °C (**Fig. 2E–H**). These parallel shifts indicate that TE remodeling contributes meaningfully to the reorganization of the proteome during heat shock. It is noteworthy that the control group exhibits a slight yet statistically significant difference in TE (**Fig. 2C, 2G**). Other processes, including protein degradation, may influence this discrepancy.

**Figure 1.**
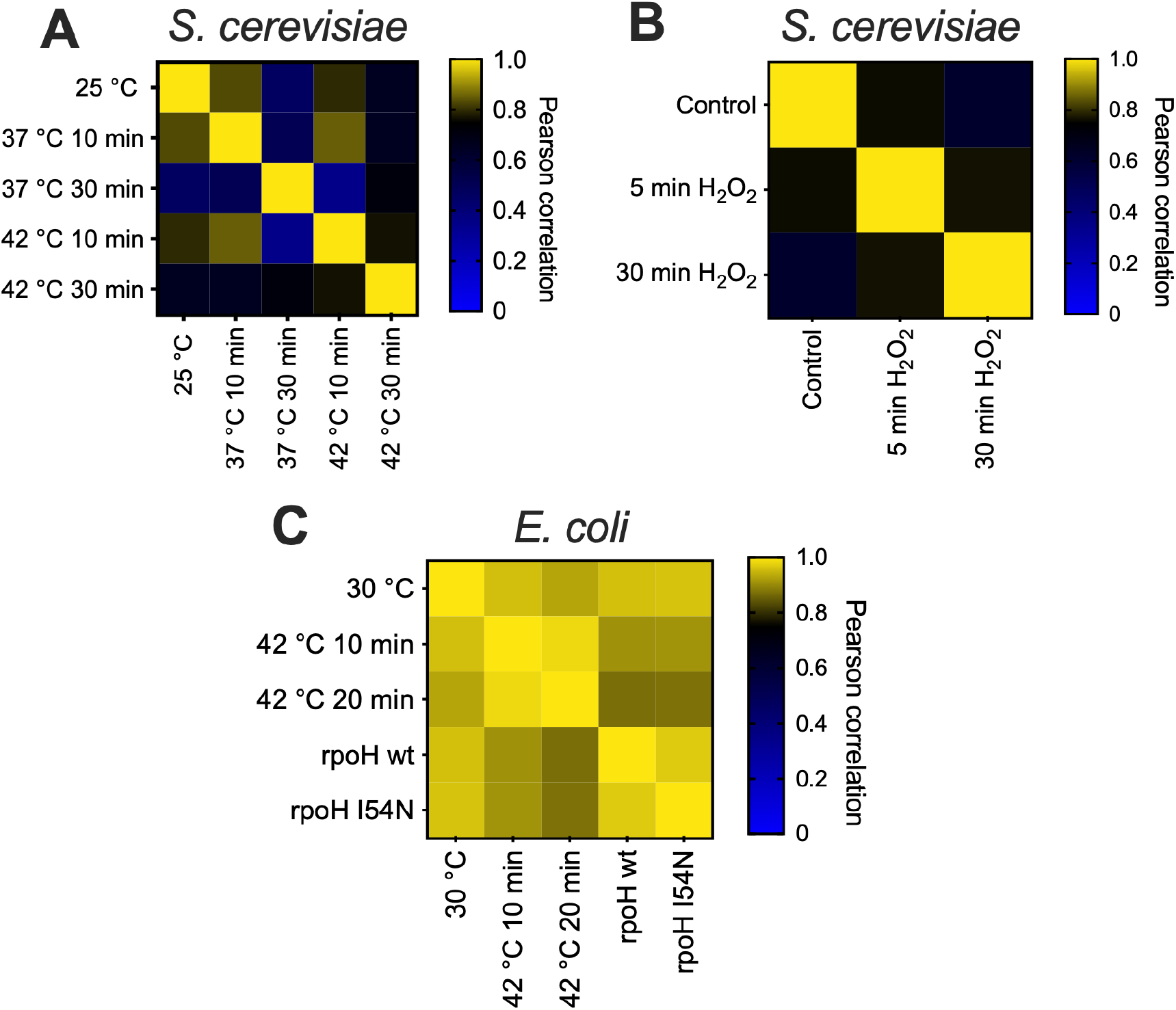
Comparison of translation efficiency under stress conditions. (A) TE values from yeast grown at 25 °C were compared with values measured after shifts to 37 °C or 42 °C. The Pearson correlation (r values) observed across different conditions indicates extensive variability in TE during heat shock. (B) Similar variability was detected under oxidative stress in yeast. (C) In contrast, TE correlations in *E. coli* remained high across different temperatures and σ32 activation, suggesting a more stable translational response than in *S. cerevisiae*.

**Figure 2.**
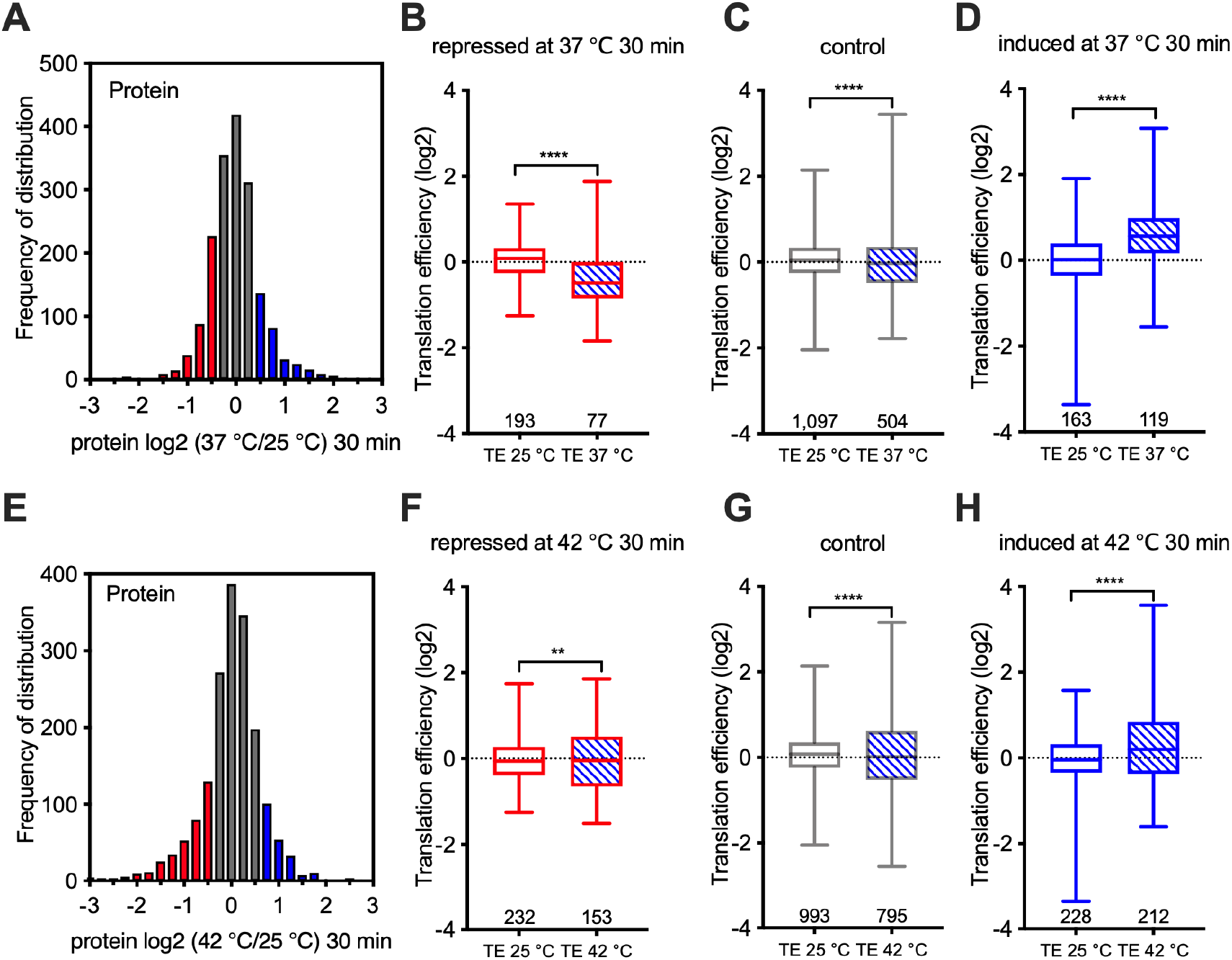
Relationship between TE changes and protein abundance during heat shock. (A)Protein abundances measured at 25 °C and after 30 min at 37 °C were grouped according to their relative changes. TE values calculated under matching conditions revealed that decreases (B) or increases (D) in protein abundance were accompanied by corresponding adjustments in TE. Proteins with a Fold Change between the two samples within the range −0.5 to 0.5 were used as control groups. The same pattern was observed under stronger heat shock at 42 °C (E-H).

Next, we assessed the changes under different heat-shock durations – specifically 10 and 30 minutes. TE values change at 10 min were examined by grouping transcripts according to their TE ratios at 37 °C relative to 25 °C (**Fig. 3A**). The transcripts exhibiting the largest early increases in TE shared specific characteristics: they tended to have shorter 5′UTRs (**Fig. 3B**) and a higher proportion of optimal codons (**Fig. 3C**). These features are generally associated with efficient translation and may provide an advantage during the initial adjustment to stress (**Razumova et al, 2025, Wang et al, 2016**). Another mechanism that can influence TE is the reprogramming of tRNA pools to regulate codon-biased translation under specific conditions, such as heat stress (**Mitchener et al., 2023**). A study from Babu’s group demonstrated that yeast adjusts the abundance of particular tRNAs to selectively enhance the translation of stress-response genes, including those involved in the heat-shock response (**Torrent et al., 2018**). The tRNA Adaptation Index (tAI) is a quantitative measure of how well a gene’s codon usage matches the availability and efficiency of the corresponding tRNAs (**dos Reis et al., 2004**). Because tRNA pools change during stress, a modified metric—the stress-adjusted tAI (s.tAI)—was developed to capture better codon–tRNA relationships under these conditions in *S. cerevisiae* (**Torrent et al., 2018**). Surprisingly, in our analysis the s.tAI values of transcripts with increased TE were lower than those of the control group (**Fig. 3D**). Instead, their TE increases were primarily associated with greater ribosome occupancy (**Fig. 3E**), with minimal alterations in mRNA abundance (**Fig. 3F**). When the heat-shock temperature was increased to 42 °C, a similar distribution of early TE changes was observed (**Fig. S1A–D**), although in this case reduced transcript abundance, coupled to high ribosome occupancy contributed more strongly to the TE increase (**Fig. S1E–F**).

**Figure 3.**
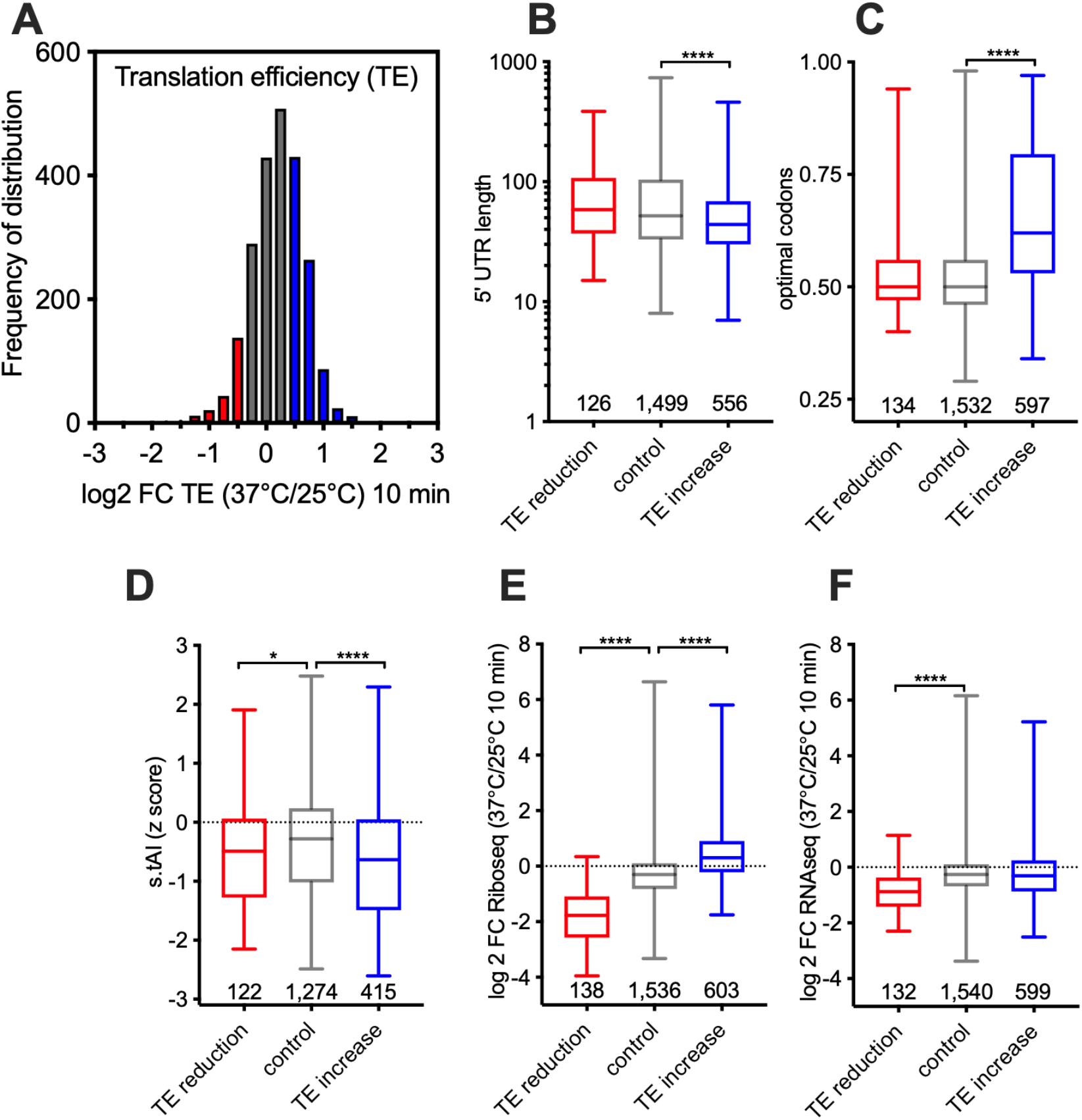
Characteristics of transcripts displaying early TE increases. After 10 min at 37 °C (A), transcripts with higher TE tended to have shorter 5 ′ UTRs and higher codon optimality (C). These TE changes were associated primarily with increased ribosome occupancy (E) rather than changes in transcript levels (F). Stress-adjusted tAI values (D) did not increase for these transcripts. Similar trends were observed under stronger heat shock Fig S1.

Although the early TE changes were substantial, they showed little predictive value for the protein abundance measured after 30 min. TE values obtained at 10 min correlated poorly (r = 0.11) with protein levels at 30 min, whereas TE values measured at 30 min correlated more strongly (r = 0.46). This indicates that the early translational response reflects a transient adjustment phase rather than the steady-state conditions that govern later protein synthesis.

To clarify how TE evolves as stress persists, we repeated the feature analysis using data collected after 30 min at 37 °C. The set of transcripts showing increased TE at this later stage differed markedly from those identified earlier (**Fig. 4A**). Furthermore, the correlation between TE increase and sequence features, such as UTR length and codon optimality composition, that were observed at 10 min under heat shock, was no longer observed after 30 min. At 30 min, transcripts with higher TE did not exhibit shorter 5′UTRs (**Fig. 4B**) or elevated optimal codon content (**Fig. 4C**), and the stress-adjusted tAI displayed a different pattern from the early response (**Fig. 4D**). Instead, TE increases at 30 min were accompanied by pronounced rises in ribosome occupancy (**Fig. 4E**) as well as higher mRNA abundance (**Fig. 4F**). The same general behavior was observed during heat shock at 42 °C (**Fig. S2**). These trends indicate that sustained TE changes do not depend on the intrinsic sequence features that influenced the early response but instead reflect broader shifts in transcript availability and ribosome access.

We found that bacteria maintain translation efficiency (TE) across different temperatures, whereas yeast does not (**Fig. 1**). It is well established that bacterial ribosomes elongate at approximately 15–20 amino acids per second (**Zhu et al., 2019**), while yeast ribosomes elongate at roughly 5 amino acids per second (**Sharma et al., 2019**). Therefore, a heat-stress exposure of the same duration (10 minutes) may affect translation differently in these organisms, potentially explaining our observations.

**Figure 4.**
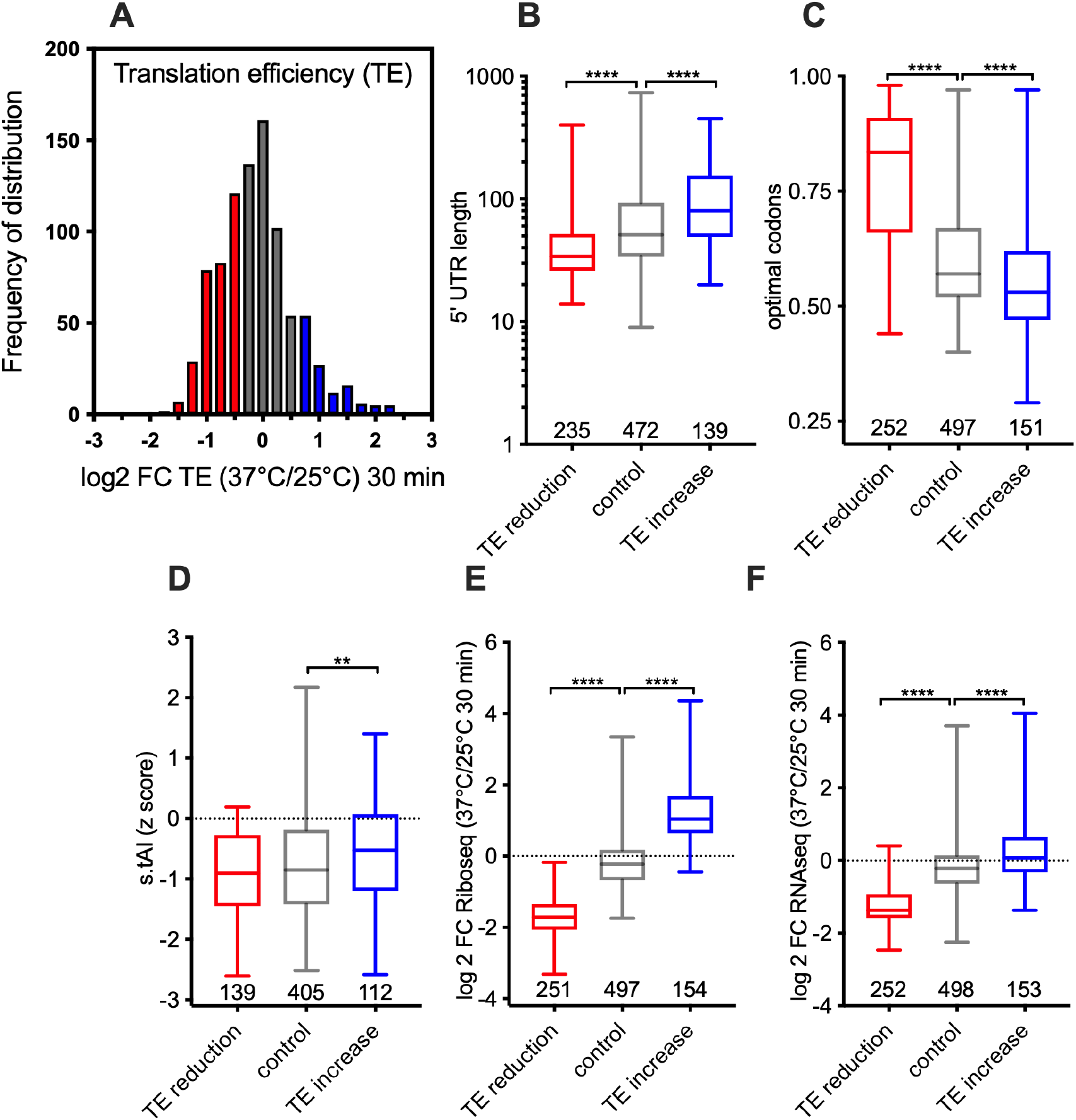
Characteristics of transcripts displaying later TE increases. After 30 min of heat shock, transcripts exhibiting higher TE differed from those identified at 10 min and lacked the same sequence features. (A) Frequency distribution of a ratio among TE 37 °C/25 °C 30 minutes. 5′UTRs (B), codon optimality (C), and stress-adjusted tAI values (D) were calculated for each group of genes. TE increases at this stage were accompanied by increases in both ribosome occupancy (E) and mRNA abundance (F).

Several mechanisms have recently been suggested to explain the temporal differences observed. Temperatures above 39 °C induce condensation of Ded1, an RNA helicase that interacts with structured 5′UTRs, and this condensation tends to sequester transcripts bearing long or structured 5′ leaders (**Iserman et al, 2020**). Such behavior would favor the translation of short 5 ′ UTR transcripts early in the response, consistent with the patterns observed at 10 min. In parallel, heat shock and other stresses promote condensation of mRNAs synthesized before the stress, while newly transcribed mRNAs remain excluded from these assemblies and available for translation (**Glauninger et al, 2025; Zedan et al, 2025**). As heat shock persists, older RNAs remain condensed and unavailable for translation, allowing free ribosomes to translate recently produced transcripts, thereby intensifying their translation regardless of features such as 5′UTR length or codon composition. This interpretation is consistent with the feature-independent TE increases observed at 30 min (**Fig. 4**).

Taken together, these findings suggest a two-phase model of translational control during heat shock. In the early phase, when cellular organization is rapidly shifting, transcripts with shorter UTRs and optimal codon usage are translated more effectively (**Fig. 3**). As the response progresses, transcriptional induction of heat-shock genes and selective sequestration of pre-existing transcripts alter the pool of ribosome-accessible mRNAs, leading to a broader reorganization of TE that is less dependent on intrinsic mRNA characteristics (**Fig 4**). Studies have shown that under heat shock, both yeast (**Rahaman et al., 2025**) and mammalian cells (**Shalgi et al., 2013**) exhibit a reduction in polysomes and a corresponding increase in monosomes, leading to a global decrease in protein synthesis. Therefore, comparisons of TE across different temperatures do not account for this shift in polysome abundance. Other phenomena, such as reprogramming tRNA (**Torrent et al., 2018)** and alternative transcription start sites (**Zan et al., 2025**), must also modulate the TE of stress response mRNAs. Through these combined mechanisms, yeast adjusts its translational landscape to support proteostasis under thermal stress.

## Acknowledgments

We thank Thais Araujo and Nicolas Meirelles for helpful discussions.

## Funding

This work was supported by Conselho Nacional de Desenvolvimento Científico e Tecnológico (CNPq), Fundação de Amparo a Pesquisa do Estado do Rio de Janeiro (FAPERJ), and Coordenação de Aperfeiçoamento de Pessoal de Nível Superior (CAPES).

### Author contributions

RLC, TD, and FLP were involved in designing and discussing the experiments. FLP was involved in drafting the paper. TD and RLC revised the paper.

## Data availability statement

The data presented in this study were obtained from public datasets and are available in Supplemental Table 1.

---

**Figure S1.**
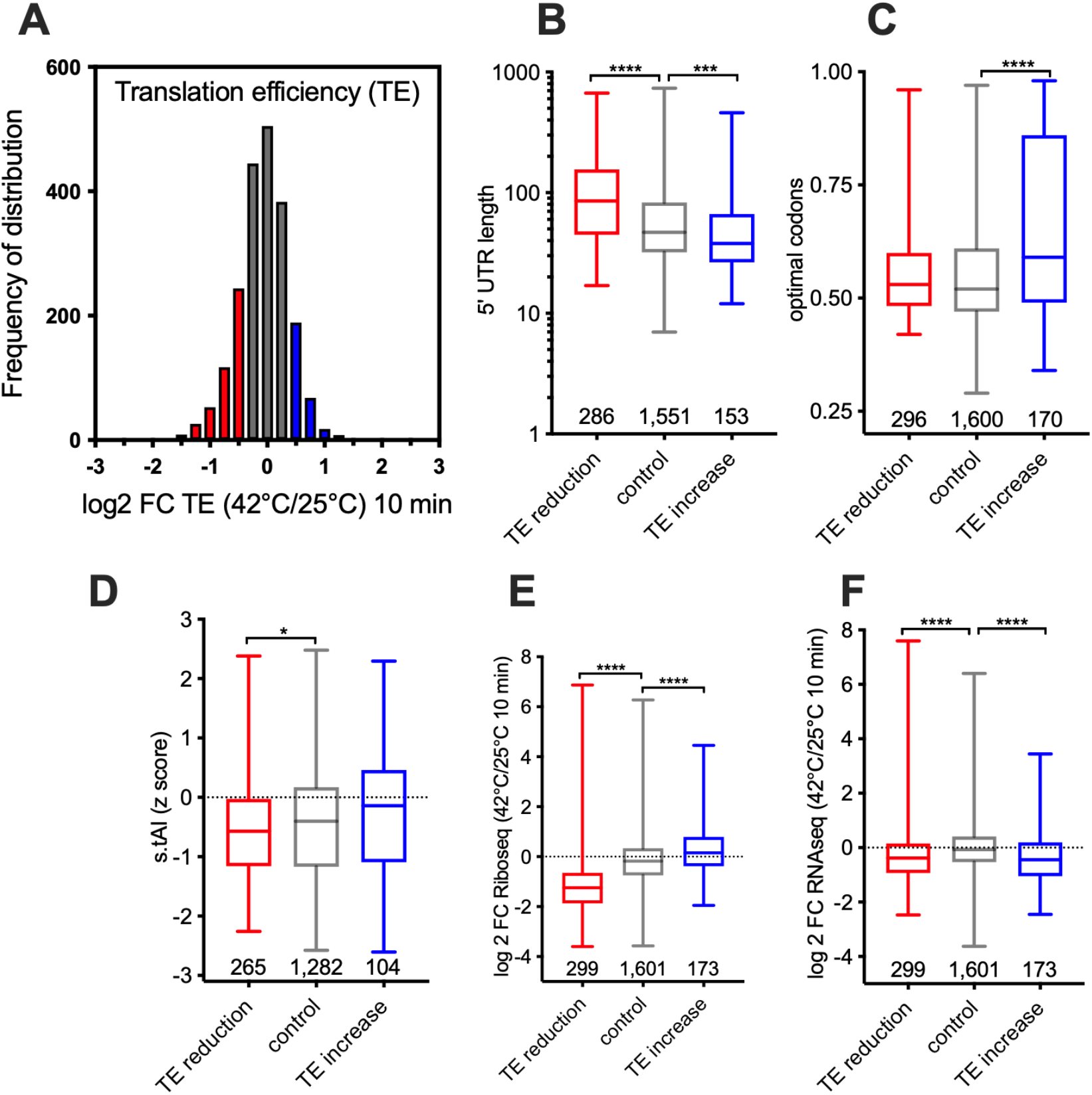
Characteristics of transcripts displaying early TE increases. (A) Frequency distribution of a ratio among TE 42 °C/25 °C 10 minutes. After 10 min at 42 °C, transcripts showing higher TE (blue symbols) tended to have short 5′UTRs (B) and higher codon optimality (C). These TE changes were primarily associated with changes in transcript levels (F) rather than with increased ribosome occupancy (E). Stress-adjusted tAI values did not increase for these transcripts (D).

**Figure S2.**
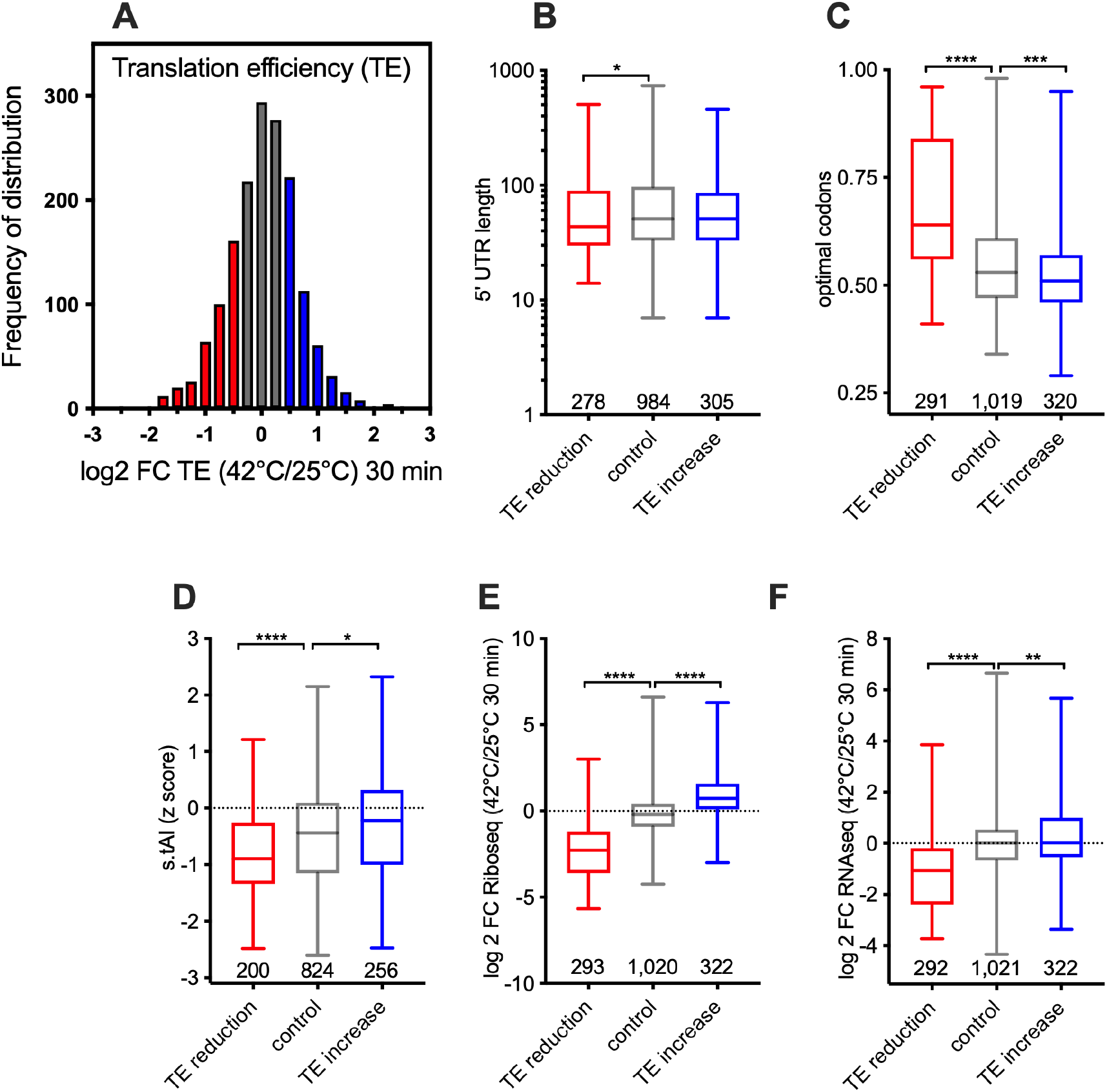
Characteristics of transcripts displaying later TE increases. After 30 min of heat shock, transcripts exhibiting higher TE differed from those identified at 10 min and lacked the same sequence features. (A) Frequency distribution of a ratio among TE 42 °C/25 °C 30 minutes. 5′UTRs (B), codon optimality (C), and stress-adjusted tAI values (D) were calculated for each group of genes. TE increases at this stage were accompanied by increases in both ribosome occupancy (E) and mRNA abundance (F).

## References

1. Desroches Altamirano C, Alberti S. Surviving the heat: the role of macromolecular assemblies in promoting cellular shutdown. Trends Biochem Sci. 2025 Jan;50(1):18–32.

2. dos Reis M, Savva R, Wernisch L. Solving the riddle of codon usage preferences: a test for translational selection. Nucleic Acids Res. 2004 Sep 24;32(17):5036–44.

3. Gerashchenko MV, Lobanov AV, Gladyshev VN. Genome-wide ribosome profiling reveals complex translational regulation in response to oxidative stress. Proc Natl Acad Sci U S A. 2012 Oct 23;109(43):17394–9.

4. Glauninger H, Bard JAM, Wong Hickernell CJ, Velez KM, Airoldi EM, Li W, Singer RH, Paul S, Fei J, Sosnick TR, Wallace EWJ, Drummond DA. Transcriptome-wide mRNP condensation precedes stress granule formation and excludes new mRNAs. bioRxiv [Preprint]. 2025 Jul 18:2024.04.15.589678.

5. Iserman C, Desroches Altamirano C, Jegers C, Friedrich U, Zarin T, Fritsch AW, Mittasch M, Domingues A, Hersemann L, Jahnel M, Richter D, Guenther UP, Hentze MW, Moses AM, Hyman AA, Kramer G, Kreysing M, Franzmann TM, Alberti S. Condensation of Ded1p Promotes a Translational Switch from Housekeeping to Stress Protein Production. Cell. 2020 May 14;181(4):818-831.e19.

6. Morgan GJ, Burkhardt DH, Kelly JW, Powers ET. Translation efficiency is maintained at elevated temperature in Escherichia coli. J Biol Chem. 2018 Jan 19;293(3):777–793.

7. Mühlhofer M, Berchtold E, Stratil CG, Csaba G, Kunold E, Bach NC, Sieber SA, Haslbeck M, Zimmer R, Buchner J. The Heat Shock Response in Yeast Maintains Protein Homeostasis by Chaperoning and Replenishing Proteins. Cell Rep. 2019 Dec 24;29(13):4593-4607.e8.

8. Ghadanian T, Iyer S, Lazzari L, Vera M. Selective Translation Under Heat Shock: Integrating HSP70 mRNA Regulation with Cellular Stress Responses in Yeast and Mammals. Mol Biol Cell. 2025 May 1;36(5):re2.

9. Li Y, Wang F, Yang J, Han Z, Chen L, Jiang W, Zhou H, Li T, Tang Z, Deng J, He X, Zha G, Hu Z, Hu Y, Wu L, Zhan C, Sun C, He Y, Xie Z. Deep generative optimization of mRNA codon sequences for enhanced mRNA translation and therapeutic efficacy. Nat Commun. 2025 Nov 12;16(1):9957.

10. Mitchener MM, Begley TJ, Dedon PC. Molecular Coping Mechanisms: Reprogramming tRNAs To Regulate Codon-Biased Translation of Stress Response Proteins. Acc Chem Res. 2023 Dec 5;56(23):3504–3514. doi:10.1021/acs.accounts.3c00572. Epub 2023 Nov 22. Erratum in: Acc Chem Res. 2024 Aug 20;57(16):2448.

11. Rahaman S, Schiffelholz N, Mittal N, Fröhlich KE, Zavolan M, Becskei A. Heat shock induces silent ribosomes and reorganizes mRNA turnover. Cell Rep. 2025 Oct 28;44(10):116447.

12. Razumova E, Makariuk A, Dontsova O, Shepelev N, Rubtsova M. Structural Features of 5’ Untranslated Region in Translational Control of Eukaryotes. Int J Mol Sci. 2025 Feb 25;26(5):1979.

13. Shalgi R, Hurt JA, Krykbaeva I, Taipale M, Lindquist S, Burge CB. Widespread regulation of translation by elongation pausing in heat shock. Mol Cell. 2013 Feb 7;49(3):439–52.

14. Sharma AK, Sormanni P, Ahmed N, Ciryam P, Friedrich UA, Kramer G, O’Brien EP. A chemical kinetic basis for measuring translation initiation and elongation rates from ribosome profiling data. PLoS Comput Biol. 2019 May 23;15(5):e1007070.

15. Torrent M, Chalancon G, de Groot NS, Wuster A, Madan Babu M. Cells alter their tRNA abundance to selectively regulate protein synthesis during stress conditions. Sci Signal. 2018 Sep 4;11(546):eaat6409.

16. Verghese J, Abrams J, Wang Y, Morano KA. Biology of the heat shock response and protein chaperones: budding yeast (Saccharomyces cerevisiae) as a model system. Microbiol Mol Biol Rev. 2012 Jun;76(2):115–58.

17. Wang X, Hou J, Quedenau C, Chen W. Pervasive isoform-specific translational regulation via alternative transcription start sites in mammals. Mol Syst Biol. 2016 Jul 18;12(7):875

18. Zedan, M., Schuerch, A.P., Heinrich, S., Garcia, P.G., Khawaja, S., and Weis, K. Newly synthesized mRNA selectively escapes translational repression following acute stress. bioRxiv, 2025 2024.04.14.589419.

19. Zhan Y, Hu Z, Lu Z, Lin Z. The patterns of alternative TSS usage explain the highly heterogeneous landscape of 5’UTR lengths in eukaryotes. NAR Genom Bioinform. 2025 Nov 11;7(4):qaf152.

20. Zhu M, Dai X. Maintenance of translational elongation rate underlies the survival of Escherichia coli during oxidative stress. Nucleic Acids Res. 2019 Aug 22;47(14):7592–7604.

